# Hippocampal BiP Overexpression Rescues Cognitive Performance and Increases REM theta in 3xTg Mouse Model of Alzheimer’s Disease

**DOI:** 10.64898/2026.03.23.713240

**Authors:** William E. Duncan, Polina Fenik, Ewa Strus, Sigrid Veasey, Nirinjini Naidoo

## Abstract

The accumulation of Aβ plaques and hyperphosphorylation of Tau neuropathologically characterize Alzheimer’s disease (AD). Synaptic dysfunction and endoplasmic reticulum (ER) stress precede overt neuropathology. ER stress is characterized by the accumulation of unfolded/misfolded proteins, which leads to activation of the adaptive signaling pathway, the unfolded protein response (UPR). Chronic or unresolved ER stress, as in disease, is maladaptive and triggers the integrated stress response (ISR). We hypothesize that targeted attenuation of ISR activation would mitigate the early cognitive deficits and molecular pathology in the triple transgenic (3xTg) mouse model of AD. To test this hypothesis, we used an adeno-associated viral (AAV) vector to overexpress BiP, the key ER chaperone and UPR regulator, in the hippocampi of young 3xTg mice. BiP overexpression reduced phosphorylated PERK (pPERK), a marker of ISR activation, and increased synaptic proteins BDNF, PSD95, and choline acetyltransferase marker (ChAT). Hippocampal-dependent working memory, social memory, long-term spatial memory, and REM theta power were improved without changes in locomotion. BiP overexpression reduced neuroinflammation, as evidenced by a decrease in the astrocyte marker GFAP. Additionally, Aβ and Aβ42 levels were reduced in the hippocampus and cortex. Collectively, these findings indicate that modulation of ER stress via BiP overexpression ameliorates early cognitive and molecular alterations associated with AD.

## Introduction

Alzheimer’s disease (AD) is an increasingly prevalent disease within the aging community, affecting about 1 in 9 Americans aged 65+ (Kanasi et al., 2016; United Nations, 2017). AD is the most common form of dementia, accounting for 60-80% of cases (Alzheimer’s Association, 2025). Accumulation of beta amyloid (Aβ) plaques and hyperphosphorylated Tau (pTau) neuropathologically characterizes AD and correlates with neural apoptosis, neurocognitive decline, and synaptic irregularity (Hardy & Selkoe, 2002). Evidence suggests that synaptic dysfunction within the hippocampus precedes the development of overt amyloid and Tau pathology, both in human and in animal models (Jacobsen et al., 2006; Meftah & Gan, 2023).

In previous studies, we have demonstrated the induction of endoplasmic reticulum (ER) stress following sleep deprivation. Disrupted sleep perturbs ER proteostasis, triggering the Unfolded Protein Response (UPR), which in turn reduces protein translation, accelerates protein degradation, and upregulates protein chaperone synthesis (Brown et al., 2014; Naidoo et al., 2008; Naidoo et al., 2005). Sleep fragmentation, reduced slow-wave sleep, reduced sleep quality, cognitive decline, and cellular stress are risk factors for AD (Aktan Süzgün et al., 2025).

We have previously demonstrated that disrupting proteostasis fragments sleep and restoring proteostasis consolidates and improves sleep (Brown et al., 2014; Hafycz et al., 2022) suggesting a bidirectional relationship between ER proteostasis and sleep. Disrupted proteostasis is a key feature in neurodegenerative diseases, including AD. Proteostasis disruption precedes AD pathology, and restoration of proteostasis, in particular the UPR, improves amyloid pathology and learning/memory in AD mouse models (Hafycz et al., 2023). The UPR maintains proteostasis in response to protein misfolding and ensuing ER stress (Hetz et al., 2020; Ron & Walter, 2007). Activation of UPR pathways is adaptive and alleviates cellular stress. Chronic ER stress/UPR activation leads to the maladaptive integrated stress response (ISR) (Pakos‐Zebrucka et al., 2016), which is observed in AD. ISR impairs the synthesis of crucial synaptic proteins, thereby contributing to memory deficits and synaptic loss that cascade into the hallmarks of late-stage AD (Hafycz et al., 2023). We therefore hypothesized that improving proteostasis via neuronal overexpression of the ER chaperone, binding immunoglobulin protein (BiP) would attenuate ISR activation and improve cognitive function in triple transgenic (3xTg) mice. To test this hypothesis, we expressed BiP in 3xTg hippocampi using an adeno-associated viral vector (AAV) based approach and assessed behavioral, electrophysiological, and molecular outcomes. Hippocampal BiP overexpression reduced ER stress, improved cognition, increased synaptic plasticity markers and reduced Aβ42 in 3xTg mice.

## Results

### Hippocampal BiP Overexpression Decreased ER Stress and Improved Cognitive Performance

To determine whether hippocampal BiP overexpression restores proteostasis and cognitive function, we delivered an AAV-CaMKII-BiP vector to the hippocampi of adult male and female 3xTg mice. An AAV-CaMKII-mCherry vector was used as a control (Figure 1A). BiP overexpression was confirmed with immunofluorescence (IF) and western blotting. BiP expressing 3xTg mice displayed increased BiP in the CA1 (p< 0.05) (Figure 1B-B’) and in the entire hippocampus when compared with controls (Figure1C).

**Fig. 1:**
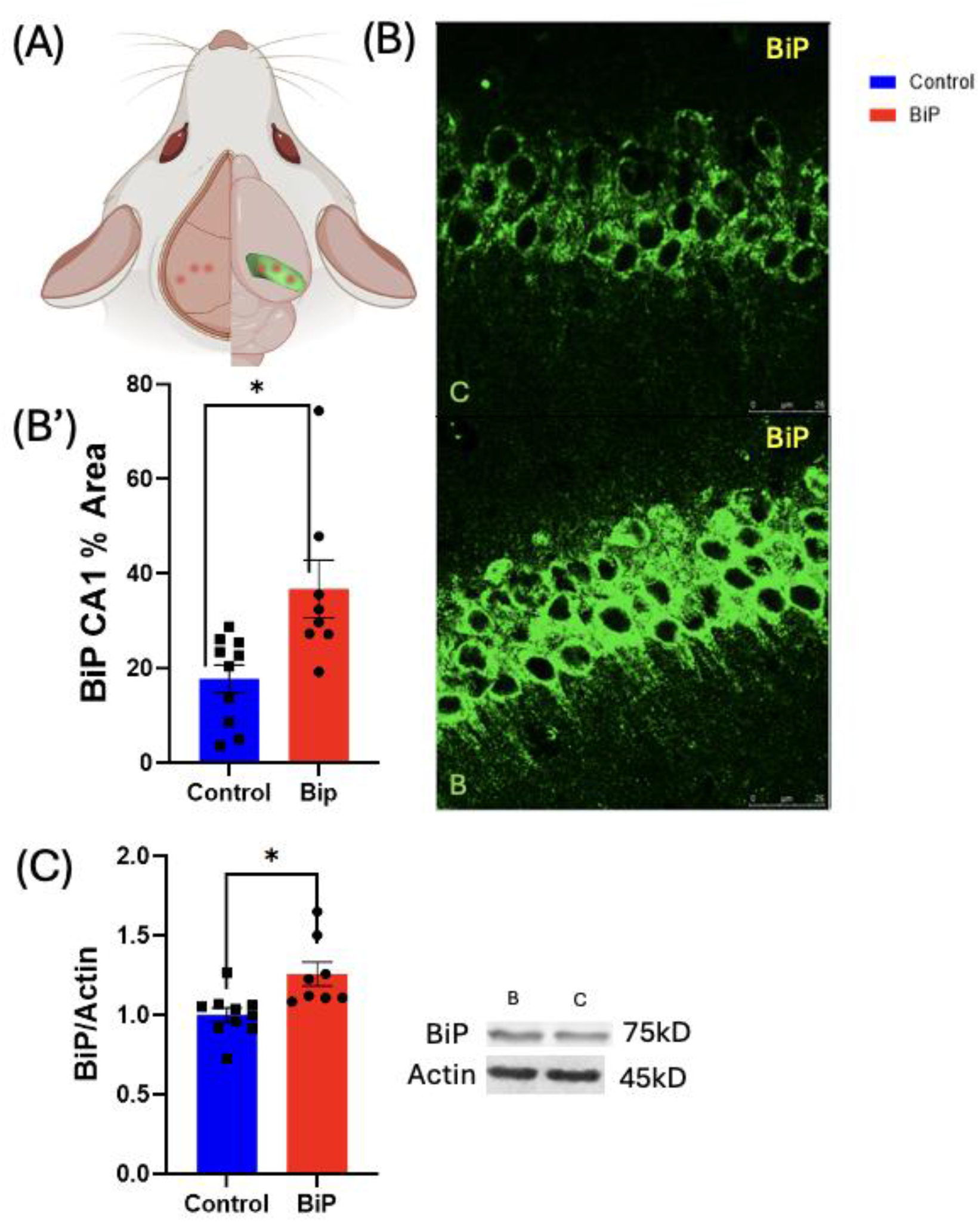
Hippocampal overexpression of BiP in 3xTg mice: A: An Illustration of the hippocampal injection created with BioRender.com: The location of the injection site on the skull and a dorsal view of the approximate injection sites into the hippocampi. B: Representative images of CA1 BiP staining. B’: unisex analysis of BiP % area of the CA1 C: unisex analysis of BiP/Actin hippocampus westerns. groups were analyzed with an unpaired t-test (n=8-10) *p<0.05. (control = C, BiP= B)

Having confirmed viral vector targeting and BiP overexpression we assessed activation of the ER stress sensor, PERK (protein kinase R-like endoplasmic reticulum kinase) that regulates protein translation. Phosphorylation of PERK results in inhibition of global protein translation. IF demonstrated that hippocampal phosphorylated PERK (pPERK) levels were significantly reduced in BiP-expressing mice compared to control mice (p<0.05) (Figure 2A-A’). This reduction was not significant in sex distinguished groups of BiP-expressing mice when compared with control mice (Figure 2A,A’’).

**Fig. 2:**
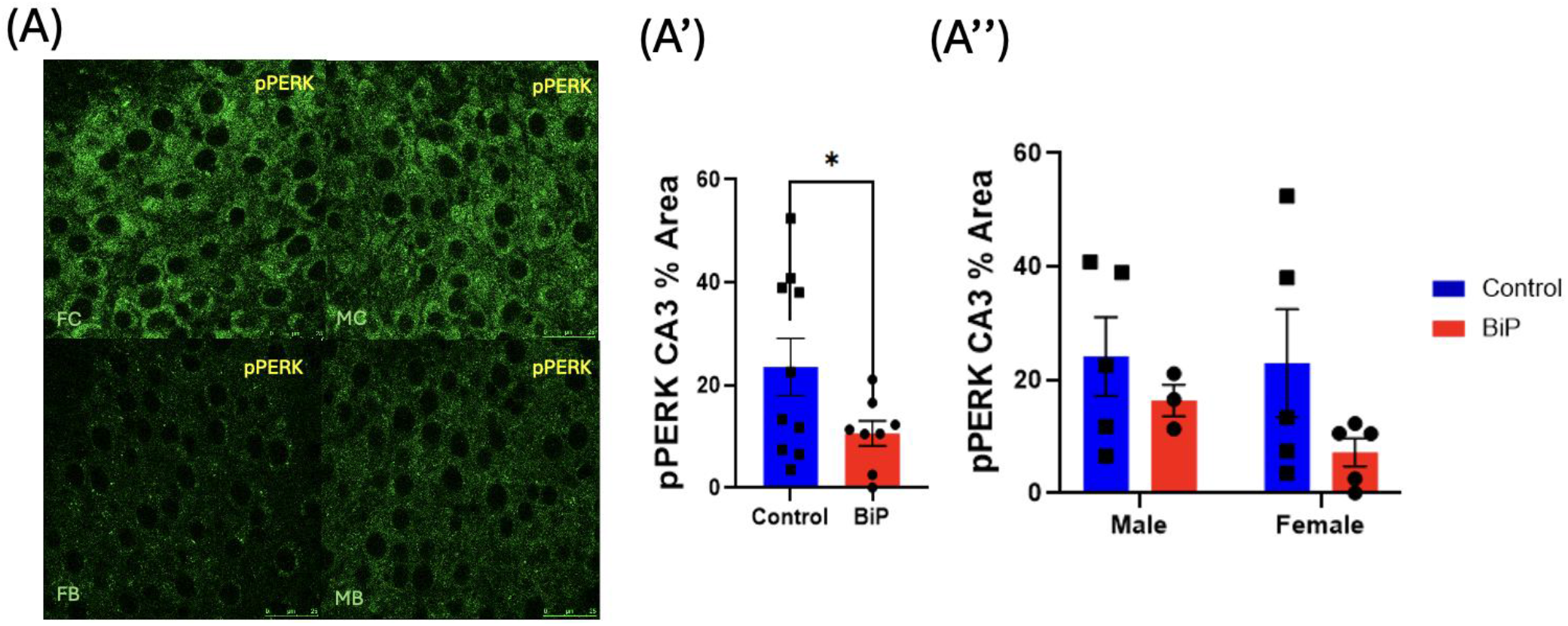
BiP Reduces PERK activation in the hippocampus: A:Representative images of CA3 p-PERK staining. A’: unisex analysis of p-PERK %area CA3 hippocampus. A’’: analysis of p-PERK %area CA3 hippocampus with sex distinguished groups. Unisex groups were analyzed with a unpaired t-test (n=8-10), *p<0.05. (Female control = FC, Male control = MC, Female BiP= FB, Male BiP= MB).

Memory deficits are a central phenotype of AD (Tarawneh & Holtzman, 2012). Many studies have shown previously that memory deficits are linked to increased ER stress and PERK activation (Ma et al., 2013; Ricobaraza et al., 2009). To determine if hippocampal BiP overexpression and reduced PERK activation was sufficient to improve learning and memory we assessed cognitive performance using the spatial object recognition (SOR), Y-maze, open field, and 3-chamber tests. BiP-expressing mice exhibited significantly improved memory and learning compared with control mice across all cognitive tests.

The SOR test was used to assess hippocampal-dependent spatial learning. Performance in the SOR test was assessed based on preference for the displaced object relative to the stationary object (Chuluun et al., 2020; Haettig et al., 2011). BiP-expressing mice displayed a significant improvement in spatial learning when compared to the control mice (p<0.05) (Figure 3A-3A’). However, when assessing sex dependent memory, male BiP-expressing mice displayed a significant improvement over their control counterparts (p< 0.01) (Figure 3A’’), whereas the increase in place preference was not significant in female mice.

**Fig. 3:**
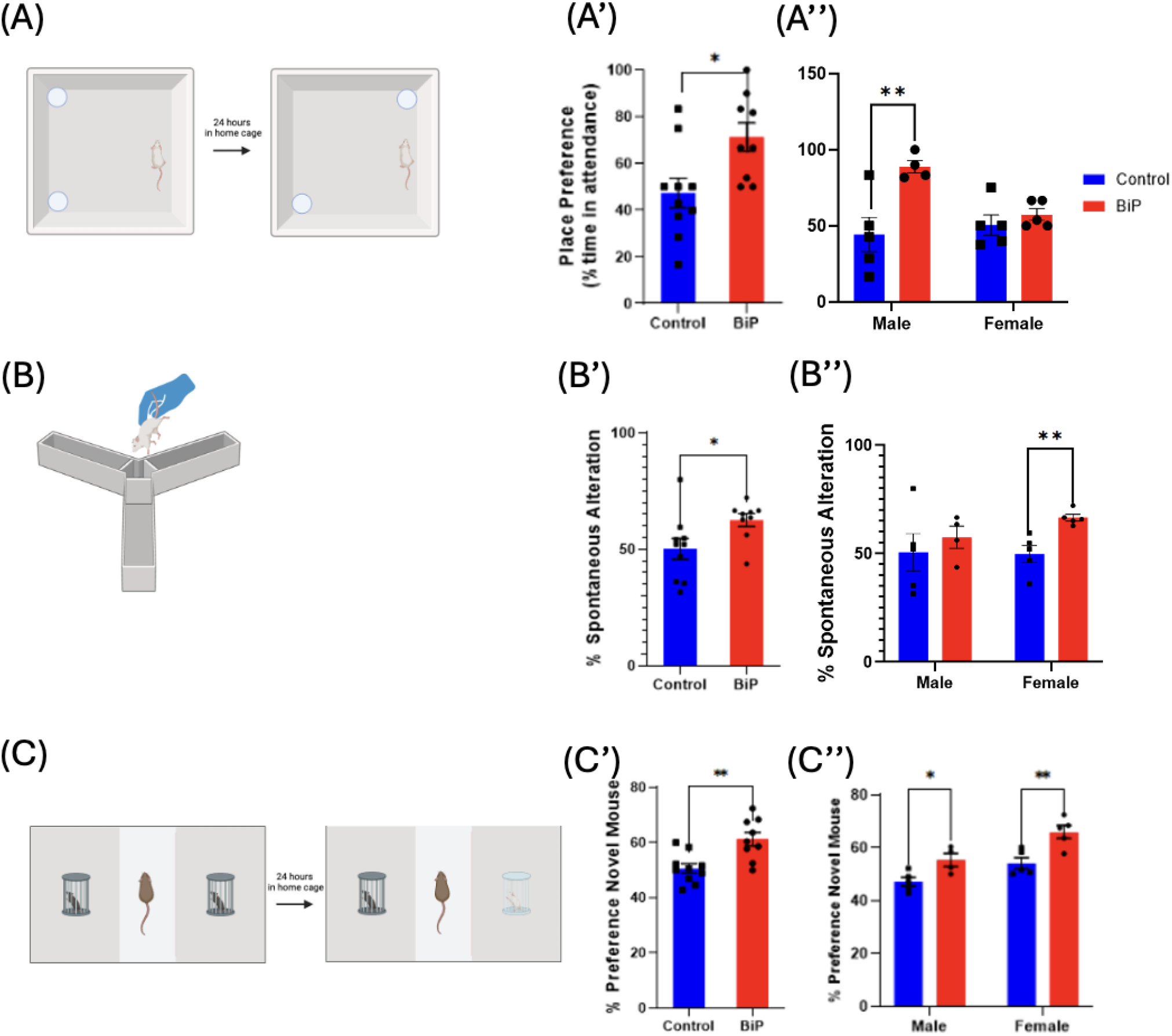
BiP overexpression improves working, social, and spatial memory: A: schematic of the SOR test. A’: SOR test with combined BiP expressing male and female groups and control male and female groups. A’’: SOR test with sex distinguished groups. B: a schematic of the Y-maze. B’: Y-maze with combined unisex groups B’’: Y-maze test with sex distinguished groups C: a schematic of the three-chamber test C’: 3-chamber with combined unisex groups C’’: 3-chamber test with sex-distinguished groups. Unisex groups were analyzed with an unpaired t-test (n=8-10), and sex distinguished graphs were analyzed with two-way ANOVA with post hoc multiple comparisons (n=3-5), *p<0.05, **p<0.01.

The spontaneous alternation Y-maze is used to test hippocampal-dependent spatial working memory (Figure 3B). Spatial working memory performance was evaluated based on spontaneous alternation behavior, expressed as the proportion of consecutive alternations relative to total arm entries. BiP-expressing mice displayed a significant improvement in spatial working memory when compared to the control mice (p<0.05) (Figure 3B’). However, when assessing sex dependent memory, female BiP-expressing mice displayed significant improvement over their control counterparts (p< 0.01) (Figure 3B’’), whereas the male mice displayed no significant difference.

The 3-chamber test is used to determine hippocampal-dependent social memory (Figure 3C). Social memory performance was evaluated based on preference for interaction with a novel mouse relative to a familiar mouse. BiP-expressing mice displayed a significant improvement in social memory when compared to the control mice (p<0.01) (Figure 3C’). Both female (p< 0.01) and male (p< 0.05) BiP-expressing mice displayed a significant improvement in social memory over their control counterparts (Figure 3C’’).

The open field test was used to assess spontaneous locomotor activity and anxiety like behavior. BiP-expressing mice and control mice exhibited no significant difference in criteria assessed for the open field maze (Distance travelled (m), mean speed (m/s), time spent on periphery/center) (see supplemental Figure 1A-C’).

### BiP Overexpression Increases Synaptic Plasticity and Memory Markers

Having observed improved cognitive performance with BiP overexpression we next examined whether increased hippocampal BiP had any effect on synaptic plasticity and memory markers. Protein levels of brain-derived neurotrophic factor (BDNF), a regulator of protein synthesis dependent memory (Lu et al., 2008; Miranda et al., 2019), the synaptic scaffolding protein, postsynaptic density 95 (PSD95), essential for synaptic stability (Caly et al., 2021), and the choline acetyl transferase marker (ChAT) which is associated with hippocampal structure and memory span (Zhu et al., 2018) were assessed by westerns and IF respectively. BiP-expressing mice displayed a significant increase in ChAT immunostaining when compared with control mice (p<0.05)(Figure 4A-A’). Analyzing ChAT levels with sex as a variable, we discovered that the increase in male BiP-expressing mice was significant when compared to their control counterparts (p<0.05) (Figure 4A,A’’). BiP-expressing mice also displayed a significant increase in BDNF levels when compared with control mice (p<0.05) (Figure 4B). Analyzing BDNF levels with sex as a variable, we discovered that the increase in female BiP-expressing mice was significant when compared to female control mice (p<0.05) (Figure 4B’). All BiP-expressing mice displayed a significant increase in PSD95 levels when compared with the control mice (p<0.05) (Figure 4C). There was no significant difference between male and female BiP-expressing mice (Figure 4C’).

**Fig. 4:**
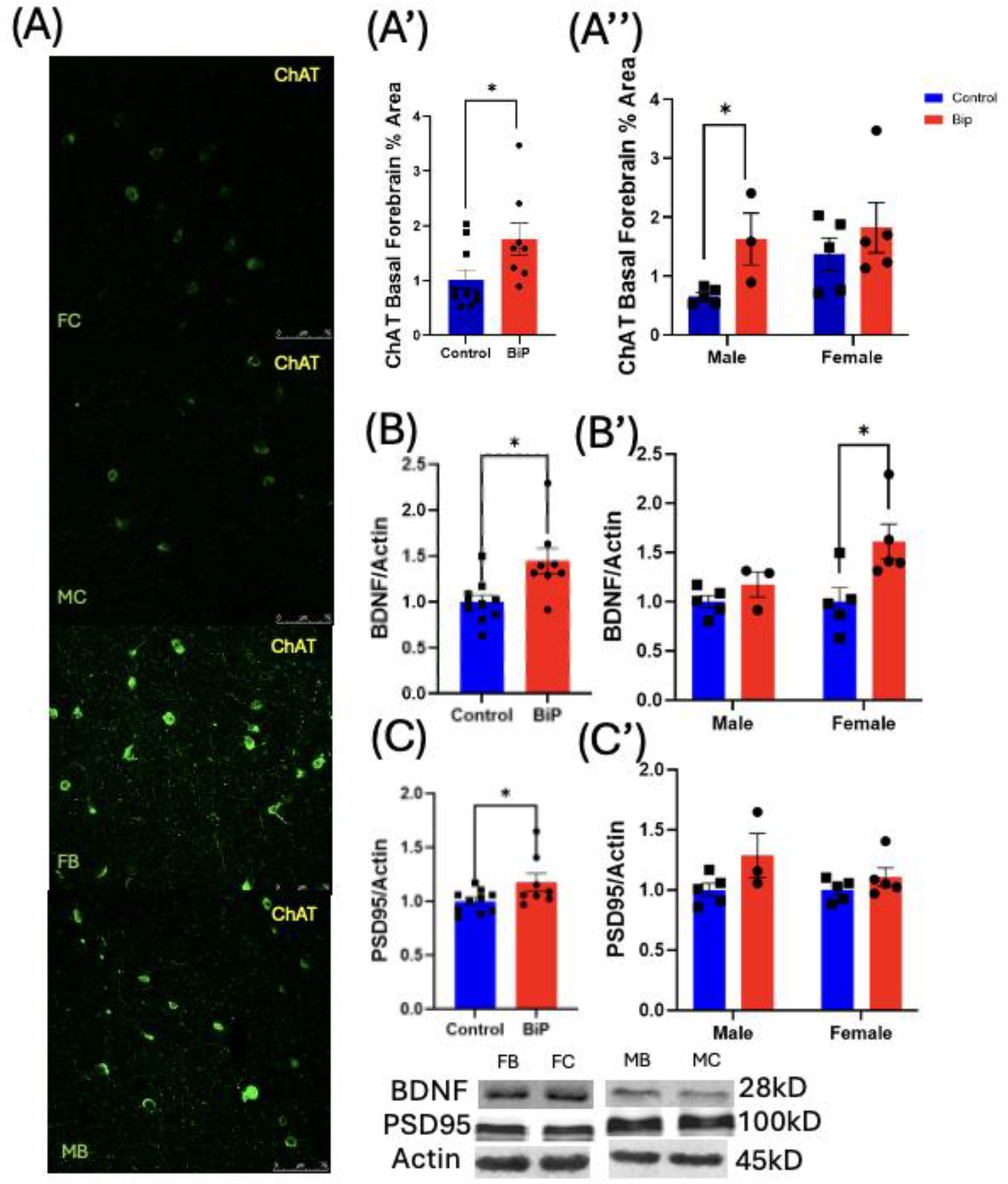
BiP increases synaptic plasticity markers: A: Representative images of basal forebrain ChAT staining. A’: unisex analysis of ChAT %area basal forebrain A’’: analysis of ChAT %area basal forebrain with sex distinguished groups B: unisex analysis of BDNF/Actin hippocampus westerns B’: analysis of BDNF/Actin hippocampus westerns with sex distinguished groups C: unisex analysis of PSD95/Actin hippocampus westerns C’: analysis of PSD95/Actin hippocampus westerns with sex distinguished groups. Unisex groups were analyzed with a unpaired t-test (n=8-10), and sex distinguished graphs were analyzed with a two-way ANOVA with post hoc multiple comparisons (n=3-5), *p<0.05. (Female control = FC, Male control = MC, Female BiP= FB, Male BiP= MB)

### BiP Overexpression Increased REM Theta Power

As sleep is known to promote memory consolidation (Lutz et al., 2026) and previous data from our group indicated that reducing ER stress with chaperone supplementation improved sleep in aged animals (Brown et al., 2014; Hafycz et al., 2022) we wanted to know whether BiP overexpression in the 3xTg mice altered sleep thereby also impacting cognition. We used Electroencephalographic (EEG) analysis to assess sleep and wake over 24 hours and found no significant differences in bout number and time spent in non-rapid eye movement (NREM), rapid eye movement (REM), or wakefulness between groups (Figure 5A-B’’). Furthermore, there were no significant differences wake or NREM bout duration (Figure 5C-C’). However, REM bout duration was significantly increased in control mice when compared to BiP mice (p<0.05) (Figure 5C’’). Having observed no major changes in sleep parameters between BiP overexpressing and control mice we decided to examine REM theta as hippocampal theta activity has been shown to increase synaptic LTP (Booth & Poe, 2006; Lutz et al., 2026; Poe et al., 2000). Spectral analyses revealed that BiP-expressing mice exhibited a significant increase in REM theta power compared to control mice (p<0.05) (Figure 5D-D’) suggesting an increase in LTP.

**Fig. 5:**
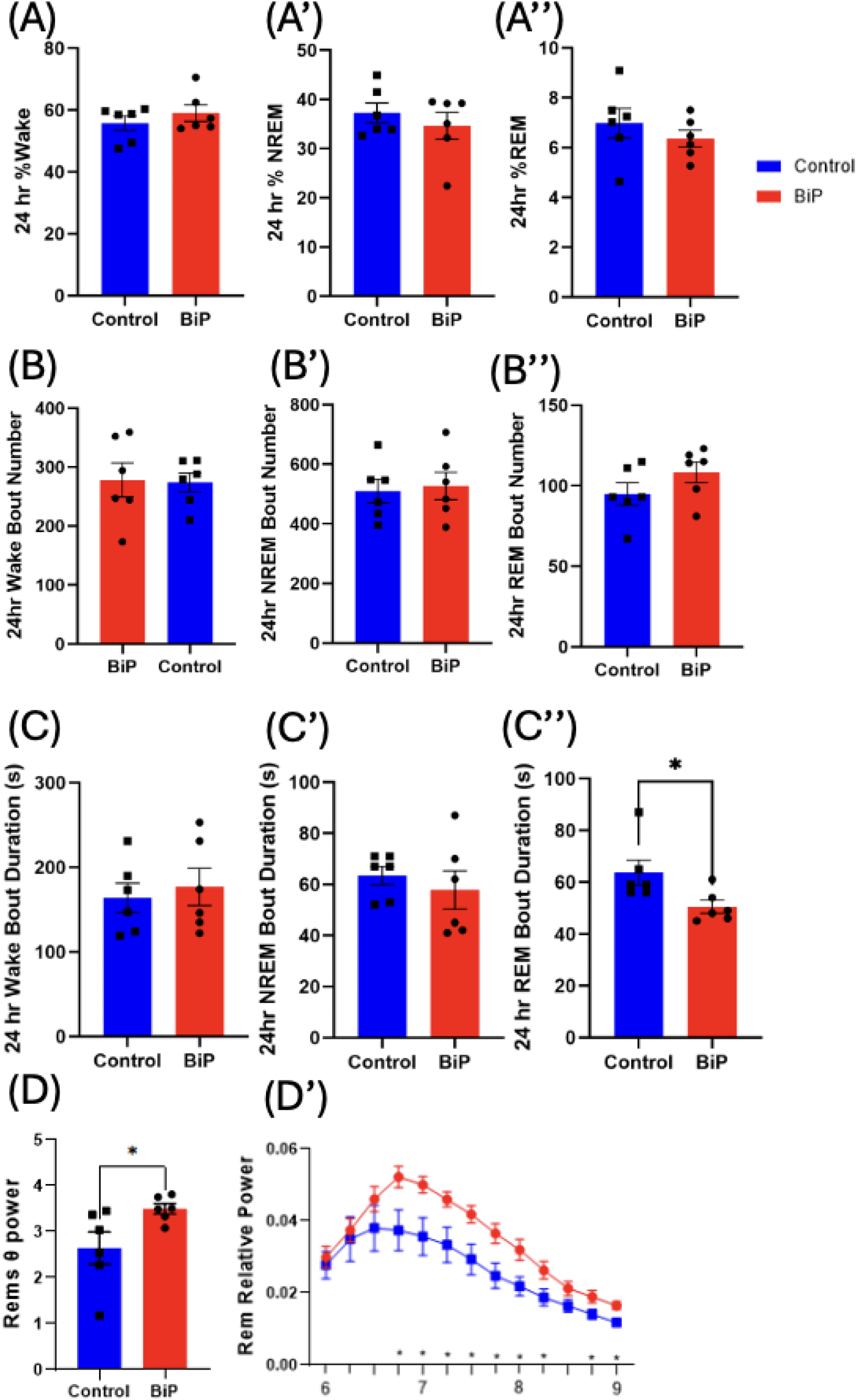
BiP Increased REM theta power: A: Unisex analysis of 24 hour % Wake. A’: Unisex analysis of 24 hour % NREM. A’’: Unisex analysis of 24 hour % REM. B: 24 hour Wake bout number. B’: 24 hour NREM bout number. B’’: Unisex analysis of 24 hour REM bout number. C: 24 hour Wake bout duration (s). C’: 24 hour NREM bout duration (s). C’’: 24 hour REM bout duration (s). D: unisex analysis of REM theta power D’: unisex analysis of REM relative power Unisex bar graphs were analyzed with an unpaired t-test (n=6), and REM Relative Power was analyzed with multiple unpaired t-test comparisons (n=6), *p<0.05.

### BiP Overexpression Reduced Neuroinflammation

Sustained neuroinflammation is a feature of AD pathology. There is evidence suggesting that astrocytes are reactivated by microglia and play an essential role in neuroinflammatory and neurodegenerative processes in AD (Liddelow et al., 2017; Rothhammer et al., 2018). To determine whether BiP overexpression had any effect on neuroinflammation, we assessed astrocyte levels using glial fibrillary acidic protein (GFAP) immunostaining. We found that control mice displayed significantly higher astrocyte levels when compared with BiP-expressing mice (p<0.0001) (Figure 6A-A’). GFAP was significantly higher in both female (p<0.05) and male (p<0.01) control mice in comparison to BiP-expressing mice (Figure 6A,A’’).

**Fig. 6:**
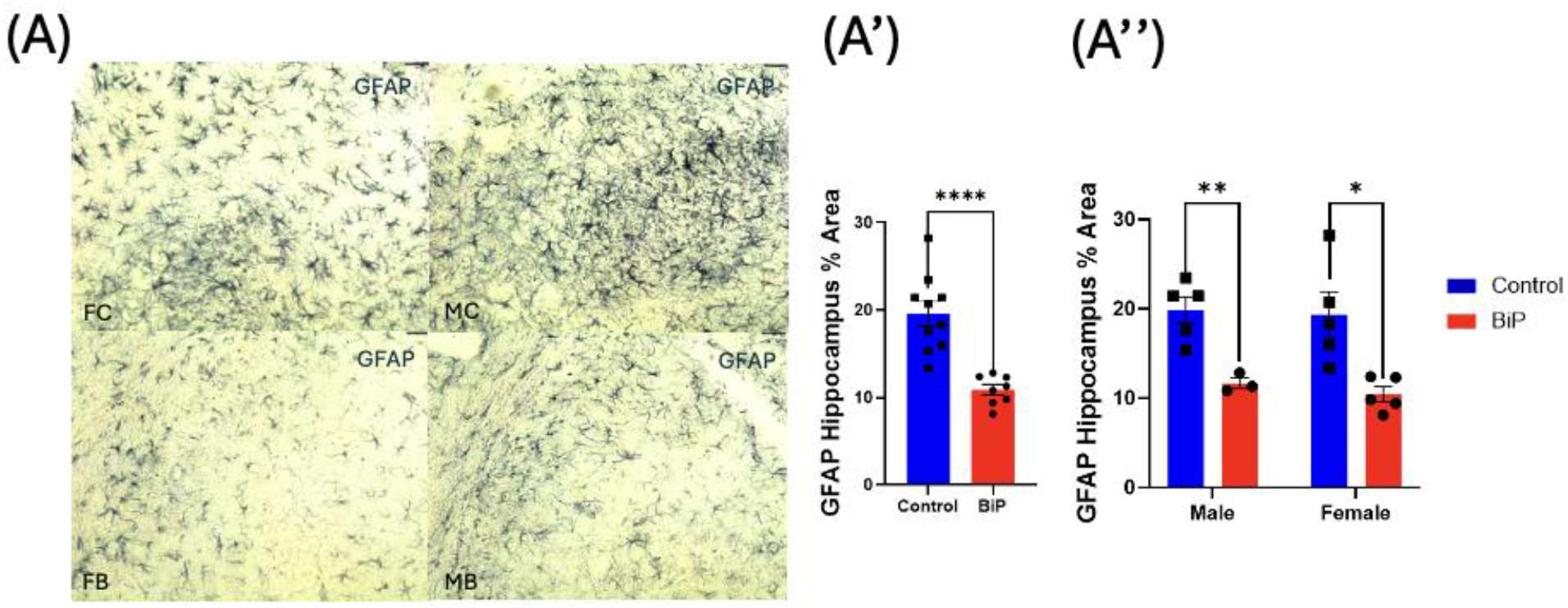
BiP decreases astrocyte levels: A: Representative images of hippocampal GFAP staining. A’: unisex analysis of GFAP %area hippocampus. A’’: analysis of GFAP %area hippocampus with sex distinguished groups. Unisex groups were analyzed with a unpaired t-test (n=8-10), and sex distinguished graphs were analyzed with a two-way ANOVA with post hoc multiple comparisons (n=3-5), *p<0.05,**p<0.01,****p< 0.0001. (Female control = FC, Male control = MC, Female BiP= FB, Male BiP= MB).

### BiP Overexpression Reduced Alzheimer’s Molecular Pathology

To determine whether overexpressing BiP in the hippocampus of 3xTg mice affected Alzheimer’s pathology, we examined phosphorylated Tau and Aβ levels in hippocampal and cortical tissue of these mice. Western blot analysis of Tau phosphorylation indicated no significant difference between BiP-expressing and control mice (see supplemental Figure 2 A-A’).

Examination of tissue stained with the 6-E10 antibody for Aβ42 revealed a few amyloid plaques within the caudal hippocampus, but not sufficient to quantify differences between groups reliably. Moreover, this staining showed increased intracellular Aβ within the subiculum of the caudal hippocampus of the control mice compared to the BiP expressing mice (p<0.05) (Figure 7A-A’). Female control mice (p<0.05) exhibited a significant increase in intracellular Aβ when compared to their BiP-expressing counterpart (Figure 7A’’). Intracellular Aβ trended lower in male BiP-expressing mice compared to the male control mice; however, it was not significant (p=0.06) (Figure 7A’’)

**Fig 7:**
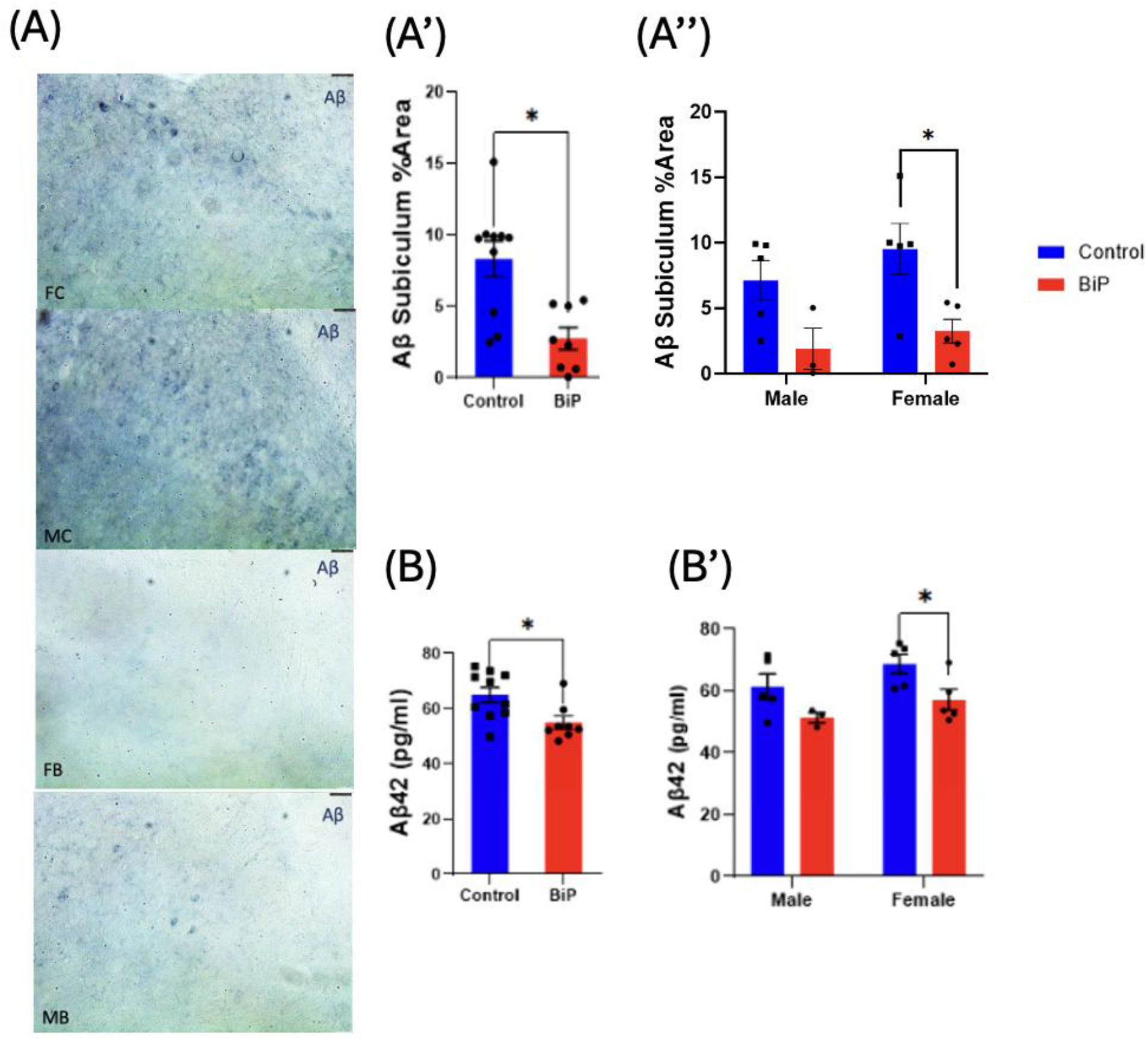
BiP Reduced Aβ and Aβ42: A: representative image of intracellular Aβ in -3.08 AP from the bregma hippocampus subiculum of each group A’: unisex analysis of intracellular Aβ hippocampus subiculum A’’: analysis of intracellular Aβ hippocampus subiculum histology with sex distinguished groups B: unisex analysis of Aβ42 cortex B’: analysis of Aβ42 cortex ELISA with sex distinguished groups. Unisex groups were analyzed with an unpaired t-test (n=8-10), and sex distinguished graphs were analyzed with two-way ANOVA with post hoc multiple comparisons (n=3-5), *p<0.05. (Female control = FC, Male control = MC, Female BiP= FB, Male BiP= MB).

Enzyme-linked immunosorbent assay (ELISA) was used to assess Aβ42 levels in cortical tissue. Control sample lysates displayed higher Aβ42 levels when compared to lysates from BiP-expressing cortices (p<0.05) (Figure 7B). Female BiP-expressing mice exhibited significantly lower Aβ42 when compared to their control counterparts (p<0.05). Aβ42 levels trended lower in male BiP-expressing mice compared to the male control mice; however, it was not significant (p=0.08) (Figure 7B’).

## Discussion

This study demonstrates that hippocampal BiP overexpression in 3xTg mice mitigates AD-associated cognitive deficits, including impairments in spatial, working, and social memory. These improvements indicate a restoration of hippocampal and cortical function (Bevins & Besheer, 2006; Cavoy & Delacour, 1993; Hafycz et al., 2022; Kraeuter et al., 2019; Sakamoto & Yashima, 2022). Importantly, the intervention did not alter locomotor activity, thereby removing activity levels as a confounding variable within the behavioral assessments.

Hippocampal BiP overexpression also increased REM theta power, a neurophysiological marker implicated in memory consolidation. Reduced REM theta power is an early indicator of changes in sleep architecture or spectral power; while this has been reported in 6-month-old 5xFAD and Tg2576 AD mouse models when compared to wildtype, it has not previously been observed in 3xTg mice. Additionally, changes in REM theta power, wakefulness, or NREM/REM duration are typically absent in 3xTg mice even at 18 months of age (Kent et al., 2018). The observed increase in REM theta power in BiP-expressing mice, therefore, suggests a beneficial modulation of sleep-related memory processes. Increased REM theta power is associated with preservation of cholinergic neurons and synaptic integrity in the hippocampus, as corroborated by our findings of increased ChAT, BDNF, and PSD95 levels in BiP-expressing mice (Gu & Yakel, 2022). Furthermore, decreases in REM theta power in humans have been associated with greater amyloid deposition (André et al., 2023).

In parallel with behavioral and electrophysiological improvements, BiP overexpression attenuated cellular stress pathways associated with AD pathology. Inhibition of the ISR via targeted BiP overexpression represents the one potential mechanism underlying the observed behavioral rescue. The ISR is a critical contributor to AD progression (Hafycz et al., 2023). Increased PERK activation within the UPR cascades to inhibition of pro-synaptic plasticity proteins and global protein translation while promoting apoptosis, ultimately contributing to synaptic failure and memory loss (Ma et al., 2013).

Consistent with these mechanisms, control mice exhibited elevated hippocampal levels of p-PERK, indicative of ISR activation. Meanwhile, reduction of p-PERK permits restoration of the translational capacity required for synaptic maintenance. Accordingly, BiP overexpression promoted BDNF and PSD95 expression, proteins essential for synaptic growth, scaffolding, and NMDA receptor regulation (Kang & Schuman, 1996; Sheng & Kim, 2011). BDNF binds to TrkB receptors on neurons, which activates downstream signaling pathways that increase expression and promote synaptic localization of PSD95 (Yoshii & Constantine-Paton, 2007). These findings are consistent with our previous work demonstrating that BiP overexpression enhances UPR signaling via IRE1/XBP1, thereby increasing BDNF levels and reducing Aβ42 accumulation in the APP_NLGF_ AD mouse model (Hafycz et al., 2023).

Beyond neuronal restoration, BiP overexpression also significantly attenuated astrogliosis, as evidenced by reduced GFAP levels. Since activation of the PERK pathway is known to influence the morphology and number of astrocytes (Smith et al., 2020), these findings suggest that BiP overexpression modulates neuroinflammation via reduced activation of PERK.

UPR signaling through the PERK pathway has also been directly linked to amyloidogenic APP processing (Devi & Ohno, 2014). At 6 months of age, 3xTg mice begin accumulating intracellular Aβ and forming Aβ plaques within the caudal subiculum and in cortical layer V (Belfiore et al., 2019; Oddo et al., 2003). In the present study, 6 month old control mice displayed an increased intracellular Aβ within the caudal subiculum, whereas 6 month old BiP-expressing mice exhibited reduced intracellular Aβ. These findings suggest that modulation of ER stress alters amyloidogenic processing at early disease stages.

Aβ42 levels were significantly reduced in the cortex of the BiP-expressing groups, indicating that hippocampal BiP overexpression may exert distal effects on cortical amyloid pathology. One potential mechanism involves disrupting the amyloid cascade by reducing hippocampal hyperexcitability and downstream cortical projections (Flores-Martínez & Peña-Ortega, 2017). Evidence indicates that pTau impairs neuronal excitability and memory consolidation in AD, preceding the development of tau tangles (Busche, 2019). It is also important to note that no difference in pTau was detected, consistent with prior reports indicating that Tau pathology emerges later in 3xTg mice. Whether sustained BiP overexpression influences Tau pathology at later stages remains to be determined (Musiek & Holtzman, 2015).

The sexual dimorphism observed in this study, as evidenced by both behavioral and molecular outcomes following hippocampal BiP overexpression, may reflect hormonal influences, particularly estrogen. Additionally, these sex distinctions could be influenced by intrinsic sex-specific differences in hippocampal circuitry and proteostatic regulation. Consistent with these sex-dependent behavioral outcomes, previous studies have shown that males exhibit more pronounced impairments in spatial and reference memory (Stevens & Brown, 2015).

Female 3xTg mice have been reported to depend on estrogen’s protective effects on cholinergic neurons and on overall AD molecular pathology (Gibbs & Aggarwal, 1998). While no studies have demonstrated a greater decline in estrogen levels in 3xTg females than in controls, loss of estrogen has been shown to be more detrimental to cellular pathology in 3xTg mice than in controls (Carroll et al., 2007). Furthermore, In AD, estrogen-related deficits manifest primarily as impaired estrogen receptor function rather than premature estrogen depletion. Chronic ER stress disrupts estrogen receptor function and downstream neuroprotective pathways, resulting in a state of functional estrogen resistance (Fan & Jordan, 2022; Ishunina et al., 2007). Further research should be conducted in early, middle, and aged 3xTg mice to determine whether estrogen receptor function is impaired and at what stage of AD progression. Estrogen is known to enhance BDNF expression and synaptic plasticity. The impaired estrogen signaling in female control mice, which then may be rescued in 3xTg BiP-expressing females, may underlie the significant improvement in spatial working memory observed in female BiP-expressing mice during the Y-maze. Sex differences in amyloid processing were observed, with females showing a more pronounced cortical Aβ42 levels which was then reduced with BiP, consistent with prior reports (Carroll et al., 2010).

The improved performance in SOR, which demands spatial encoding and consolidation, may rely more heavily on cholinergic modulation. Although previous literature reports greater cholinergic neuron swelling in female 3xTg mice than in males at later ages (13–15 months) (Perez et al., 2011), these findings are not directly comparable to the early time points examined in the present study. To our knowledge, no studies in AD mouse models have directly demonstrated earlier cholinergic dysfunction or structural loss in males relative to females at young ages, including 3 months, when behavioral tests were performed, and 6 months, when the 3xTg mice were perfused. Nevertheless, recent work demonstrates sex-dependent modulation of cholinergic signaling and amyloid pathology in AD, suggesting that cholinergic circuits are differentially regulated by sex hormones early in disease progression (German‐Castelan et al., 2024). This supports the possibility that male and female cholinergic systems diverge in vulnerability and functional regulation before overt neuronal loss becomes detectable. Additionally, during early stages of AD, including mild cognitive impairment, compensatory upregulation of ChAT in female humans has been reported which may be a mechanism to counteract emerging cognitive deficits (Dekosky et al., 2002). This upregulation of ChAT may contribute to the early cholinergic changes observed in male, but not female mice.

Together, these findings suggest that BiP-mediated improvements in proteostasis and network function engage partially distinct molecular and circuit-level pathways in males and female 3xTg mice, resulting in task-specific cognitive benefits despite a shared intervention. However, a limitation of the present study must be considered when analyzing this sexual dimorphism being the reduced sample size in certain experimental groups. Specifically, the number of male BiP-expressing mice was lower than anticipated due to post-surgical mortality, which reduced statistical power for sex-stratified analyses. In addition, EEG analyses were limited by a smaller sample size resulting from inconsistent signal quality during recording. Furthermore, 3xTg male mice have been reported to have inconsistent phenotypic expression (Duncan et al., 2022). Although these factors may have reduced sensitivity to detect sex-specific effects in some measures, the consistency of behavioral and molecular trends across independent assays supports the robustness of the overall findings.

Targeted modulation of proteostasis within the hippocampus confers functional protection against early AD-associated cognitive and molecular pathology in 3xTg mice. BiP overexpression effectively alleviated maladaptive ER stress, reduced ISR activation, enhanced synaptic protein expression, reduced neuroinflammation, and improved hippocampal-dependent memory performance. These effects were accompanied by increased REM theta power and reductions in Aβ and Aβ42, supporting a link between ER stress modulation, sleep-associated neural activity, and early amyloid pathology. Although AD pathology remained at an early stage in this model (Clark et al., 2015; Stevens & Brown, 2015), the present findings indicate that early intervention targeting ER stress may modify disease trajectory and warrant further investigation in aged 3xTg mice.

## Methods/Materials

### Mice

Animal experiments were conducted in accordance with the guidelines of the University of Pennsylvania Institutional Animal Care and Use Committee. Mice were maintained as previously described (Chellappa et al., 2019; Naidoo et al., 2008). Mice were housed at 23°C on a 12:12 h light/dark cycle and had ad libitum access to food and water. For this study, male and female 3xTg mice (n=10/group) were obtained from the National Institute of Aging. Male and female mice (n=5/group) were stereotaxically injected with AAV-BiP or the mCherry control vector.

### Stereotaxic Injection Surgery

Stereotaxic hippocampal injections were performed in accordance with the guidelines of the University of Pennsylvania Institutional Animal Care and Use Committee and with proper anesthesia, analgesia, and sterility as described in (Hafycz et al., 2022). Injections delivered an AAV-BiP vector to half of the mice (Vector Biolabs; AAV5-CamKIIa-GRP78; titer 4.8 × 1012 GC/ml) and an mCherry control vector to the other half of the mice (Addgene; pAAV5-CaMKIIa-mCherry; titer 2.3 × 10−13 GC/ml) into the hippocampi of the mice. Six burr holes were drilled for bilateral hippocampal injections, with coordinates as follows: AP -2.1 mm,-2.5mm ML ±2 mm, ±2 mm, ±1.5 mm DV-1.7 mm,−2.0mm. Using a 1 μL Hamilton syringe, 50 nanoliters of virus were injected into each burr hole. Mice were observed for 3 days of recovery. Mice were injected at 8 weeks of age with behavioral testing was conducted four weeks after surgery to allow sufficient viral expression.

### Electrode implantation, sleep recording, and analysis

Electroencephalogram surgeries were performed as previously described, with minor adjustments (Chellappa et al., 2019; Naidoo et al., 2018). Briefly, mice were anesthetized with isoflurane. Four EEG electrodes and two EMG electrodes were implanted and secured with dental acrylic. After one week of postoperative recovery, mice were individually housed, connected to EEG cables, and acclimated for three days prior to the start of recordings. Recordings took place as previously described (Naidoo et al., 2018). Recordings started at ZT0 lights on (07:00 a.m.) and continued for two consecutive days. The first 24 h of recording served as the sleep recordings. Data were scored with SleepSign Analysis Software (version 3.0, Kissei), and spectral data were analyzed as previously described (Franken et al., 2001; Hasan et al., 2012; Lim et al., 2013; Naidoo et al., 2018).

### Cognitive Tests

#### Spatial Object Recognition (SOR)

The SOR test is a well-established hippocampal-dependent spatial memory test (Bevins & Besheer, 2006; Cavoy & Delacour, 1993; Hafycz et al., 2022). SOR was conducted as previously described, with a training phase followed by a test phase 24 hours later (Hafycz et al., 2022). Preference index calculations were performed as previously described to assess how well the mice distinguished between the moved and unmoved objects (Lanzillotta et al., 2024)

#### Y-Maze

The Y-Maze test was performed as previously described (Kraeuter et al., 2019). Briefly, a single mouse was placed in the center of the apparatus and allowed to move freely through the maze for 5 min. Each individual arm entry and the order in which the entries occurred were recorded. After testing, the number of alternations (three separate arm sequential arm entries) was counted and presented as a percentage.

#### Open Field

Locomotor activity was measured in the Open Field test as previously described (Seibenhener & Wooten, 2015). Mice were placed in the center of the open field and allowed to roam for 10 minutes. The trials were recorded and analyzed using a video tracking system ANY-maze (version 7.62, Stoelting). Distance travelled (m), speed, mobility, and time spent in different zones of the arena were calculated.

#### Three Chamber

Mice underwent the Three Chamber Social Approach Test as previously described (Kaidanovich-Beilin et al., 2011). Briefly, the experimental mouse is allowed to interact with the “familiar” mice, which are placed in a cylinder at the end of the maze. 24 hours following this, a novel mouse is placed in one cylinder and a familiar mouse in the other cylinder, and the experimental mouse was allowed 10 minutes to roam and interact with both mice.

#### Immunohistochemical Assay

Postfixed half-brain coronal sections were sliced at 40 μm using a cryostat as previously described (Hafycz et al., 2022; Zhu et al., 2007). Every other section was placed in 24-well plates containing cryoprotectant for free-floating immunohistochemistry staining and stored at −20 °C, as previously described (Naidoo et al., 2008; Naidoo et al., 2018). For all markers, we compared n = 3–5 in each of the four groups. Primary antibodies are as follows: beta-amyloid 6E10 clone (1:1000, Biolegend), GFAP (1:1000, Cell Signaling); Secondary antibodies are as follows: Biotin-SP Donkey Anti-Mouse IgG (1:500, Jackson ImmunoResearch)

#### Immunofluorescence (IF)

Immunofluorescence staining was performed as previously described (Hafycz et al., 2022; Naidoo et al., 2011). We compared n = 3–5 in each of the four groups. Primary antibodies are as follows: BiP/anti-KDEL (1:1000, Enzo Life Sciences), Phospho-PERK (1:200, ThermoFisher), (ChAT (1:100, Invitrogen); Secondary antibodies are as follows: Alexa Fluor 488 donkey anti-mouse IgG (1:500).

#### Quantitative analysis of Immunofluorescence Images

Images were obtained using a Leica SP5/AOBS (confocal) and Leica DM5500B (light microscopy) microscope as previously described (Hafycz et al., 2022; Nick et al., 2022; Owen et al., 2021; Zhu et al., 2016). For Confocal image acquisition, intensities, nm range, detector gain, exposure time, amplifier offset, and focal-plane depth within sections per antigen target were standardized across the sections being compared. Two sections per animal (n = 5 mice/group) were imaged. Using ImageJ software, the images were converted to 8-bit grayscale with a detection threshold standardized across images to detect the % area. The %area covered within the selected region was measured, and the average %area for each mouse was analyzed.

#### Western blot assay

Frozen brain tissue was prepared for Western blot assays as previously described (Naidoo et al., 2008; Naidoo et al., 2018). Homogenized tissue samples (20 μg protein) were separated by SDS-PAGE, and protein bands were imaged and quantified via either Odyssey scanner (LiCor) or Alpha Innotech FluorChem 8900 (IMGEN Technologies). For all markers, n=3-5 samples were compared. Primary antibodies are as follows; BiP/GRP78 (1:1000, Enzo Life Sciences);: beta-amyloid 6E10 clone (1:500, Biolegend); BDNF (Abcam 1:1000); pTau (genetex 1:500); Tau (cell signaling 1:1000); PSD95 (Thermo Fischer 1:1000); β-Actin (1:2000, Santa Cruz sc-47778). Secondary antibodies are as follows: LiCor IRDye 680RD Goat anti-Mouse (1:10,000);LiCor IRDye 800RD Goat anti-Mouse (1:10,00 0); LiCor IRDye 800RD Goat anti-Rabbit (1:10,000); and Odyssey IRDye 680 Goat anti-Rabbit (1:10,000). Goat anti-Mouse IgG HRP (1:2000).

#### Enzyme-linked immunosorbent assay

To quantify Aβ42 in cortical tissue, an ELISA was performed according to manufacturer instructions (KMB3441; Invitrogen) and as previously described in (Yang et al., 2023).

## Data and Statistical analyses

Data are presented as the average ± standard error of the mean (SEM) of the sample size. We did not hypothesize there would be a sex-specific effect, therefore, we pooled all samples and found BiP overexpression samples were significantly different from controls. To specifically rule out a sex-specific effect, we stratified our samples by sex and found the significance was diminished likely due to the low sample size. The direction of the effect was maintained in both sexes; however, we cannot entirely rule out sex as a modifying factor. Statistical analyses were performed using GraphPad Prism (version 11.0.0, GraphPad Software). Unless otherwise specified, two-way ANOVA was used to determine interaction effects, with post hoc multiple comparisons. p < 0.05 was the threshold for determining statistical significance.

## Author Contributions

NN and SV designed the study; WD performed the experiments; WD drafted the initial manuscript; WD and PF performed IHC, ES performed WB, PF analyzed sleep data. NN revised the manuscript; NN provided the resources for the project; all authors approved of the final draft of the manuscript.

## Data Availability Statement

The raw data supporting the conclusions in this manuscript will be made available upon request.

