## supplemental figures 1 and 2 for "Hippocampal BiP Overexpression Rescues Cognitive Performance and Increases REM theta in 3xTg Mouse Model of Alzheimer’s Disease"

Supplementary Materials

Figure Legends

FigS1: Overexpression of BiP had no effect on anxiety or locomotion within the open field test: A: unisex analysis of distance travelled within the open field test A’: analysis of distance travelled within the open field test with sex distinguished groups B: unisex analysis of mean speed (meters per second) within the open field test B’: analysis of mean speed (meters per second) within the open field test with sex distinguished groups. C: unisex analysis of time spent in the periphery of the open field test C’: analysis of time spent in the periphery of the open field test with sex distinguished groups. Unisex groups were analyzed with an unpaired t-test (n=8-10), and sex distinguished graphs were analyzed with two-way ANOVA with post hoc multiple comparisons (n=3-5).

FigS2: Overexpression of BiP had no effect on pTau levels: A: unisex analysis of pTau/Tau within the hippocampus B analysis of pTau/Tau within the hippocampus with sex distinguished groups. Unisex groups were analyzed with an unpaired t-test (n=8-10), and sex distinguished graphs were analyzed with two-way ANOVA with post hoc multiple comparisons (n=3-5). (Female control = FC, Male control = MC, Female BiP= FB, Male BiP= MB).

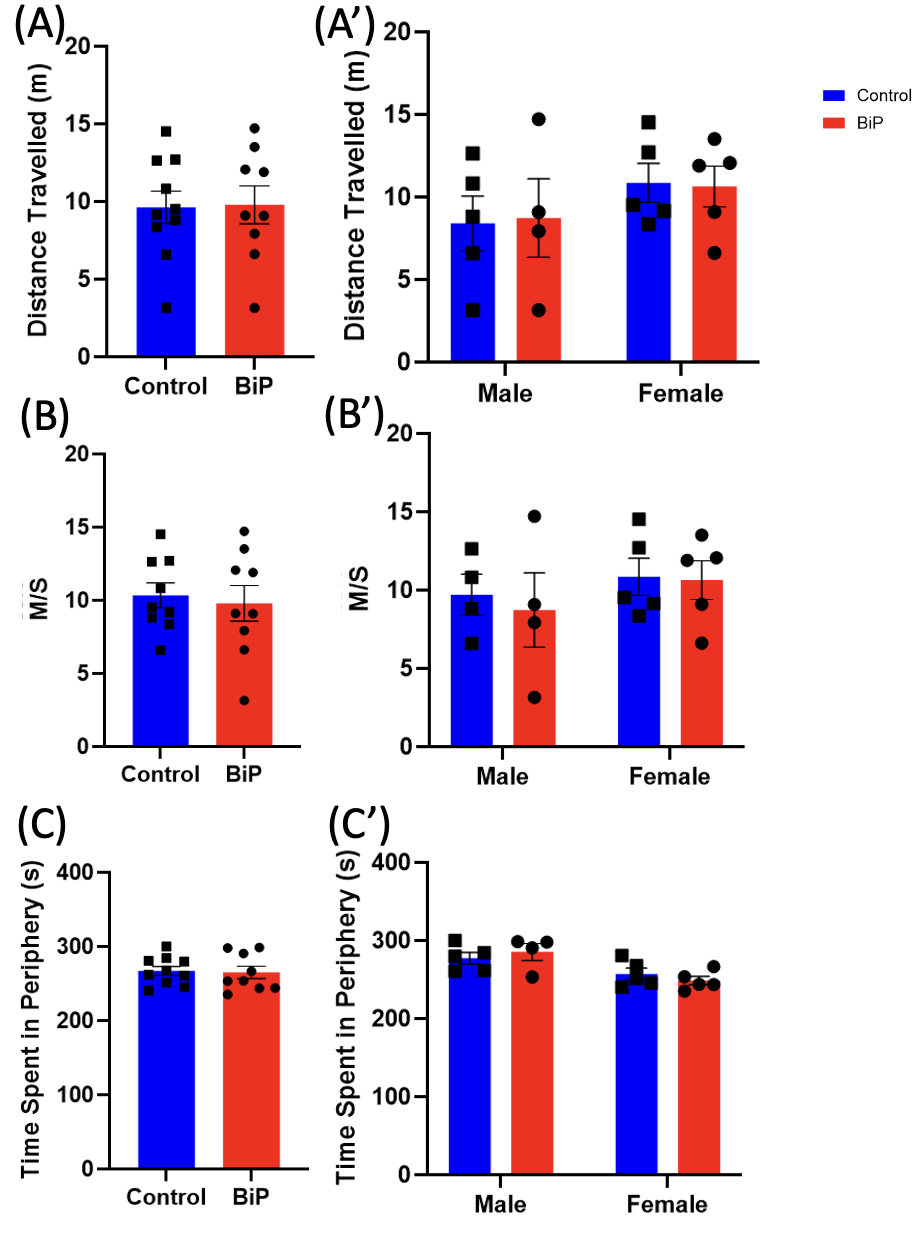

FigS1:Overexpression of BiP had no effect on anxiety or locomotion within the open field test: A: unisex analysis of distance travelled within the open field test A’: analysis of distance travelled within the open field test with sex distinguished groups B: unisex analysis of mean speed (meters per second) within the open field test B’: analysis of mean speed (meters per second) within the open field test with sex distinguished groups. C: unisex analysis of time spent in the periphery of the open field test C’: analysis of time spent in the periphery of the open field test with sex distinguished groups. Unisex groups were analyzed with an unpaired t-test (n=8-10), and sex distinguished graphs were analyzed with two-way ANOVA with post hoc multiple comparisons (n=3-5).

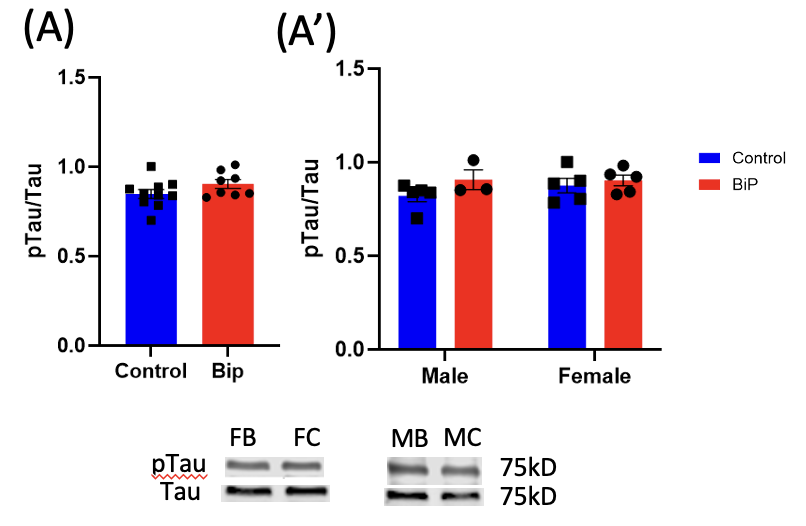

FigS2: Overexpression of BiP had no effect on pTau levels: A: unisex analysis of pTau/Tau within the hippocampus B analysis of pTau/Tau within the hippocampus with sex distinguished groups. Unisex groups were analyzed with an unpaired t-test (n=8-10), and sex distinguished graphs were analyzed with two-way ANOVA with post hoc multiple comparisons (n=3-5). (Female control = FC, Male control = MC, Female BiP= FB, Male BiP= MB).
